# Tissue accumulation and the effects of long-term dietary copper contamination on osmoregulation in the mudflat fiddler crab *Minuca rapax* (Crustacea, Ocypodidae)

**DOI:** 10.1101/2020.04.20.051508

**Authors:** MV Capparelli, JC McNamara, MG Grosell

## Abstract

We examined copper accumulation in the hemolymph, gills and hepatopancreas, and hemolymph osmolality, Na^+^ and Cl^-^ concentrations, together with gill Na^+^/K^+^-ATPase and carbonic anhydrase activities, after dietary copper delivery (0, 100 or 500 µg Cu/g) for 12 days in a fiddler crab, *Minuca rapax*. In contaminated crabs, copper concentration decreased in the hemolymph and hepatopancreas, but increased in the gills. Hemolymph osmolality and gill Na^+^/K^+^-ATPase activity increased while hemolymph [Na^+^] and [Cl^-^] and gill carbonic anhydrase activity decreased. Excretion likely accounts for the decreased hemolymph and hepatopancreas copper titers. Dietary copper clearly affects osmoregulatory ability and hemolymph Na^+^ and Cl^-^ regulation in *M. rapax*. Gill copper accumulation decreased carbonic anhydrase activity, suggesting that dietary copper affects acid-base balance. Elevated gill Na^+^/K^+^-ATPase activity appears to compensate for the ion-regulatory disturbance. These effects of dietary copper illustrate likely impacts on semi-terrestrial species that feed on metal contaminated sediments.

## Introduction

Analyses of different routes of metal uptake are essential to understanding bioaccumulation and toxicity in semi-terrestrial organisms, particularly those inhabiting environments under tidal influence. Uptake from the dissolved water phase and from ingested food may both be important routes of metal accumulation (Vitale *et al* 1999). Dissolved metals can accumulate by direct adsorption to body surfaces and by absorption across the respiratory epithelia while particulate metals can accumulate following the ingestion and digestion of food (Wang and Fisher, 1999). In crustaceans, most studies have concentrated on metals dissolved in the water phase (see review by Viarengo and Nott, 1993), and only a very few investigations have examined the effects of metal contaminated diets (Sá *et al.* 2008, Sabatini *et al.* 2009, Bordon *et al*. 2018). Our current understanding of dietary metal toxicity is inadequate, and consequently, the dietary contamination route is not usually considered in existing regulations concerning environmental contamination or in risk assessments (Borgmann *et al*. 2005, Schamphelaere *et al*. 2007).

Copper is an essential micronutrient required by all living organisms for a variety of physiological and biochemical processes. This metal is a co-factor in multiple enzymatic processes but is potentially toxic to aquatic organisms above certain levels (Martins *et al.* 2011). During exposure to elevated copper titers in the water or diet, cellular detoxification mechanisms may become saturated to a point where protein function is impaired (Grosell *et al*. 2012). In crustaceans, copper is a component of the respiratory pigment hemocyanin used in oxygen transport (Rainer and Brouwer 1993). However, while high copper titers may cause respiratory disruption in freshwater organisms, the mortality resulting from environmentally relevant waterborne copper contamination usually derives from osmoregulatory disturbances (Grosell *et al.* 2002). Copper is present in all aquatic environments, and multiple anthropic activities such as industries, agriculture and harbors discharge effluent containing elevated copper, leading to increased exposure and potential toxicity to aquatic organisms (Martins *et al.* 2011). Complexation of copper with organic and inorganic ligands as well as competition with other cations for binding and uptake pathways greatly influence copper toxicity in fresh water (Grosell *et al.* 2007).

Although decapod crustaceans are mainly aquatic, many species show varying degrees of terrestriality (Burggren and McMahon 1988). Thus, dietary contamination by heavy metals is particularly relevant in semi-terrestrial crabs where waterborne exposure is episodic and often occurs only during high tide. Nevertheless, very few studies have investigated different routes of metal contamination and chronic toxicity. Dietary metal exposure appears to affect mainly reproduction (Lauer and Bianchini 2010, Bielmyer *et al.* 2006) and energy metabolism (De Schamphelaere *et al.* 2007). In contrast, waterborne metals exert a general effect, more related to osmoregulatory disruption (Capparelli *et al*. 2016, 2017). In aquatic organisms, metal uptake generally occurs via epithelial surfaces related to gas exchange, ion absorption and secretion such as the gills of crustaceans and fish (Péqueux, 1995; Santore *et al.*, 2001). Some metals compete with other cations for binding and active uptake sites in the gills (Paquin *et al.*, 2002; Grosell *et al.*, 2007), and once absorbed, may accumulate in various tissues, particularly the gills, leading to diverse toxic effects (MacRae *et al* 1999; Grosell *et al* 2007).

In aquatic environments, copper and other non-essential trace metals exert their toxic action in a fashion synergistic with salinity variation, and gill Na^+^/K^+^-ATPase and carbonic anhydrase activities can be affected in osmoregulating crustaceans during exposure to waterborne metal contamination (Roast *et al*. 2002, Capparelli *et al*. 2017). The exact mechanisms of metal toxicity are unclear but many metals cause enzyme kinetic changes that disrupt specific metabolic systems. Na^+^/K^+^-ATPase and carbonic anhydrase activities are of special concern owing to their role in ion transport by crustacean gills, and may be particularly vulnerable to waterborne pollutants (Böttcher 1991, 1993, Kjoss et al., 2005, Bianchini *et al.* 2003).

The species used as a model for copper contamination in the present study is the semi-terrestrial, mudflat fiddler crab, *Minuca rapax*. This crab is distributed from Florida, throughout the Gulf of Mexico, the Antilles and Venezuela to the Atlantic coast of Brazil where it occurs from Pará to Santa Catarina States (Thurman *et al*. 2013), inhabiting burrows in muddy sand in estuarine mangrove environments. *Minuca rapax* is a strong euryhaline osmoregulator and maintains its hemolymph osmolality at around 780 mOsm kg^-1^ H_2_O over a wide range of environmental and experimental salinities (Thurman *et al*. 2017). Populations of this crab are affected by waterborne copper contamination, showing tissue accumulation and impaired osmotic and ionic regulatory ability (Capparelli *et al*. 2017) and metabolic and oxidative stress (Capparelli *et al*. 2019).

Fiddler crabs are common inhabitants of mangrove biotopes and are important sediment bioturbators, feeding avidly on sediment grains from which they glean organic matter, algae, micro-organisms and bacteria, which are ingested as food, together with small inorganic particles (Crane 1975, Christy 1978, Kristensen 2008). Given that *M. rapax* spends much of its time feeding on sediment particles, evaluating the impacts of long-term dietary contamination should provide important insights into copper accumulation and toxicity. We address two questions in the present study: (i) does dietary copper contamination lead to greater tissue bioaccumulation than does waterborne exposure; and (ii) does dietary copper accumulate in the gills and affect osmotic and ionic regulatory ability?

## Materials and Methods

Adult, intermolt specimens of *Minuca rapax* of either sex were collected from Virginia Key Beach (25° 44’ 28.17” N, 80° 0.8’ 50.74” W), Virginia Key, Miami, Florida and transported in plastic boxes containing sponge cubes moistened with water from the collection site to the Laboratory of Environmental Physiology and Toxicology at the University of Miami, Florida. Only non-ovigerous, intermolt crabs of carapace width greater than 10 mm were used. To acclimatize to laboratory conditions before use, the crabs were maintained unfed for three days at 25 °C, with free access to a dry surface, in plastic boxes containing water from the collection site.

An artificial diet containing 10% dry mass was prepared by mixing 2 g agar, 3 g sucrose and 5 g fish food (which contains 18% protein, 16% fiber and 2.5% crude fat, calorie content 2,300 to 2,500 kcal/kg, and 8,000, 1,000, and 50 IU/kg vitamins A, D3 and E, respectively) with solutions containing the requisite concentrations of copper chloride (100 or 500 µg/g) to give 100 mL of agar medium (Swaileh and Ezzughayyar 2000). 100 mL of each copper-containing medium was divided equally among four petri dishes (25 mL/dish), which after cooling, were kept in a refrigerator at 4 °C. During the experiments, agar cubes were cut out, adjusted to 0.15 g weight and placed in the plastic boxes containing the individual crabs. A control food source without copper chloride (0 µg/g) was prepared in exactly the same way.

The crabs were kept for 12 days (N= 10 per trial) in the laboratory during the experiments. They were sorted randomly into individual plastic boxes containing dilute seawater (25 ‰ salinity, 750 mOsm kg^-1^ H_2_O, 350 mmol Na^+^ L^-1^, 400 mmol Cl^-^ L^-1^) in one section and the 0.15-g cube of artificial food in a dry section. During the 12-day experimental period, the crabs were fed with a known amount of food to which either 0, 100 or 500 µg CuCl_2_/g agar had been added. At the end of each day, any leftover food remnants were removed, the seawater was changed and a new food cube was offered. Usually, all food was consumed within the 24-h period.

Following the 12-day period of dietary copper contamination, the crabs were cryo-anesthetized in crushed ice for 5 min each. A hemolymph sample was drawn into a 1 mL syringe using a needle inserted into the arthrodial membrane at the base of the 3rd or 4th pereiopod and frozen at -20 °C. The crabs were then killed by removing the carapace, and the anterior and posterior gill pairs and hepatopancreas were dissected from each animal (N= 10). The hemolymph and tissues were frozen at -80 °C for later use.

After thawing, hemolymph osmolality was measured in 10 µL aliquots using a vapor pressure micro-osmometer (Wescor Model 5500, Wescor Inc., Logan, UT, USA). Hemolymph [Na^+^] was measured using a Varian FS220Z atomic absorption spectrophotometer (Varian, Mulgrave, Victoria, Australia). Hemolymph [Cl^-^] was quantified employing anion chromatography (Dionex DX-120, fitted with an AS40 automated sampler, Dionex Corp., Sunnyvale, CA, USA).

After chilling briefly on ice, the pooled anterior and posterior gills from each crab were homogenized in dry ice and acetone using a buffer containing (in mmol L^−1^) imidazole 20, pH 6.8, sucrose 250, EDTA 6 and a protease inhibitor cocktail (Furriel et al., 2000). The hydrolysis of *p*-nitrophenylphosphate ditris salt (*p*NPP) (i. e., the K^+^-phosphatase activity of the Na^+^/K^+^-ATPase) by the gill homogenates was assayed continuously for 15 minutes at 25 °C by monitoring the release of the *p*-nitrophenolate ion at 410 nm spectrophotometrically (Spectramax Plus 384 microplate reader, Molecular Devices) under the standard conditions described by Furriel *et al.* (2000).

The K^+^ phosphatase activity of the Na^+^/K^+^-ATPase was assayed by adding aliquots of each sample homogenate to a reaction medium containing (in mmol L^-1^) HEPES buffer 50, pH 7.5, KCl 10, MgCl_2_ 5, *p*NPP 10 (for total K^+^-phosphatase activity) or the same medium containing 3 mmol L^−1^ ouabain, a specific inhibitor of *p*NPPase activity (for oubain insensitive activity). The K^+^ phosphatase activity in each sample was estimated from the difference between the total *p*NPPase activity and the ouabain insensitive activity.

To assay carbonic anhydrase activity, the pooled anterior and posterior gills from each crab were homogenized using a Potter homogenizer in homogenization buffer (in mmol L^−1^, mannitol 225, sucrose 75 and Trizma-base 10, pH 7.4). The homogenate was then differentially centrifuged to separate the cytoplasmic and membrane-bound isoforms. Cytoplasmic carbonic anhydrase activity was measured using an electrometric method (Henry, 1991).

Measurement of total protein in all homogenates for all assays was performed according to Bradford (1976) employing bovine serum albumin as the standard.

Copper content in the hemolymph, pooled anterior and posterior gills, and in the hepatopancreas was measured by atomic absorption spectroscopy, employing a graphite furnace (Varian FS220Z atomic absorption spectrophotometer, Mulgrave, Victoria, Australia) using certified reference material. Matrix interference in the diluted seawater medium (25 ‰S) was resolved using a solvent extraction technique (Blanchard and Grosell, 2006).

The samples were digested for 24 h in 1 N HNO_3_ (1: 10 w/v, Trace Metal Grade) at 60 °C, vortexed and centrifuged, and the supernatants were collected for analysis after appropriate dilution.

All data are expressed as mean values ± the standard error of the mean (SEM). After verifying normality of distribution and equality of variance, the physiological and biochemical parameters were analyzed using two-way (effect of copper concentration and tissue on copper accumulation) or one-way (effect of copper concentration on hemolymph osmolality, [Na^+^] and [Cl^-^], and gill Na^+^/K^+^-ATPase and carbonic anhydrase activities) analyses of variance, followed by the Student-Newman-Keuls *post-hoc* multiple comparisons procedure. Occasionally, raw data were transformed to meet normal distribution criteria. Differences were considered significant at P=0.05.

## Results and Discussion

No mortality was recorded in *M. rapax* receiving dietary copper. The crabs did not alter their behavior and they did not avoid the copper-contaminated feed, each crab consuming the entire agar cube offered daily.

The two-way analysis of variance revealed that copper accumulation was affected by both tissue and copper concentration and their interaction (19.2 < F < 98.1, P < 0.001). In the control crabs (no added Cu), [Cu] were highest in the hemolymph (518 µg Cu/g), lowest in the gills (40 µg Cu/g) and intermediate in the hepatopancreas (360 µg Cu/g) (0.001 < P < 0.023) (Figure 1). Curiously, [Cu] were lower in the hemolymph (P < 0.001) and hepatopancreas (0.017 < P < 0.001) of the crabs receiving the copper-contaminated diets, particularly 500 µg Cu/g, than in the control crabs. In the gills, [Cu] was highest in the crabs receiving 500 µg Cu/g (P < 0.001) (Figure 1).

**Fig. 1.**
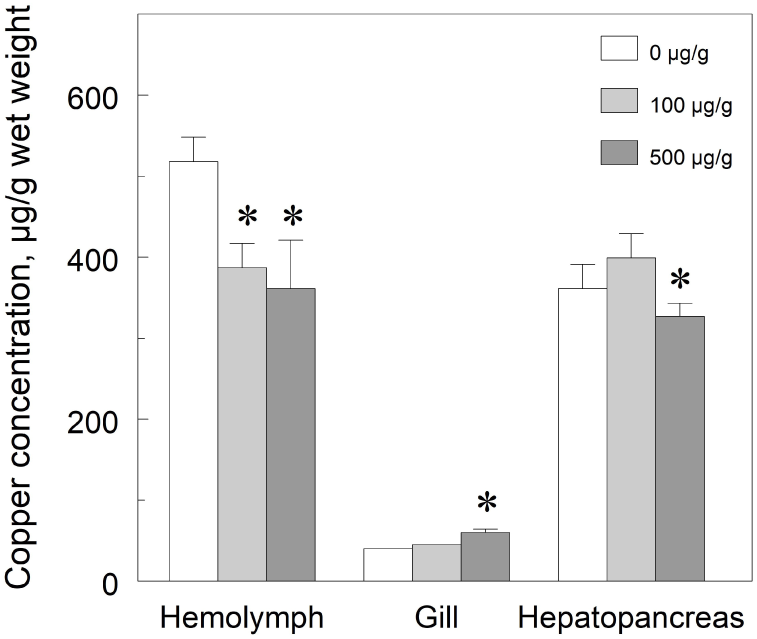
Copper concentrations in the hemolymph, pooled anterior and posterior gills, and hepatopancreas of *Minuca rapax* receiving a copper-contaminated diet (100 or 500 µg CuCl_2_/g) for 12 days. Data are the mean ± standard error (N= 10). *P≤ 0.05 compared to control crabs (0 µg CuCl_2_/g).

Hemolymph osmolality was slightly hyper-regulated (845 ± 0.9 mOsm kg^-1^ H_2_O, Δ ≈+100 mOsm kg^-1^ H_2_O) in the control crabs and strongly hyper-regulated (Δ ≈+200 mOsm kg^-1^ H_2_O) above ambient values in the copper contaminated crabs. Sodium was strongly hyper-regulated (518 ± 1.4 mmol L^-1^, Δ ≈+170 mmol L^-1^) in the control crabs although less so in the copper-contaminated crabs (Δ ≈+40 mmol L^-1^) while chloride was isocloremic (413 ± 0.9 mmol L^-1^) in the control crabs and slightly hypo-regulated (Δ ≈-35 mmol L^-1^) in the copper-contaminated crabs.

Hemolymph osmolality was higher in the crabs receiving the copper-contaminated diets (0.021 < P < 0.037) (947 mOsm kg^-1^ H_2_O) compared to the control crabs (845 mOsm kg^-1^ H_2_O). However, hemolymph sodium (P < 0.001) and chloride (0.034 < P < 0.048) concentrations were lower (≈380 mmol L^-1^) in these same crabs compared to the control crabs (518 mmol Na^+^ L^-1^, 413 mmol Cl^-^ L^-1^) (Figure 2).

**Fig. 2.**
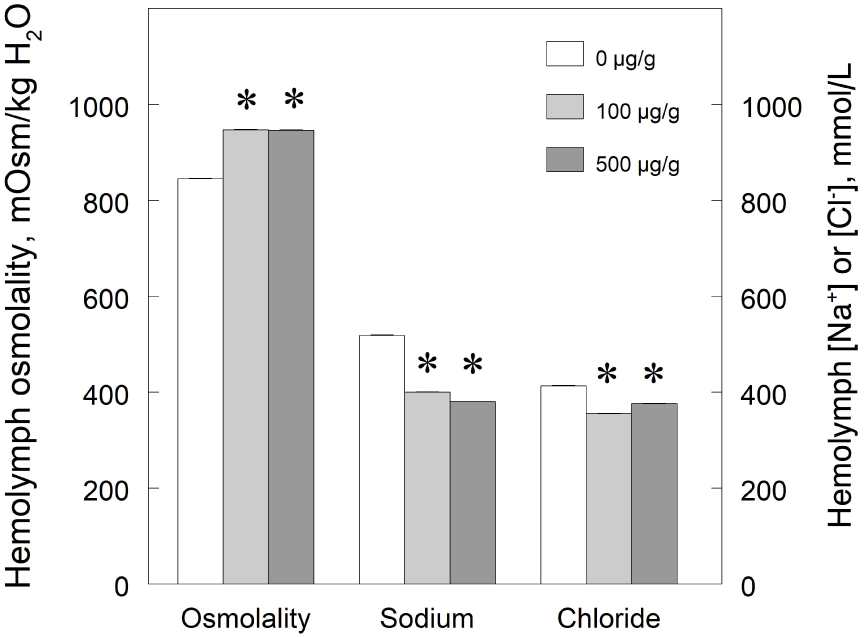
Hemolymph osmolality, sodium and chloride concentrations in *Minuca rapax* fed a copper-contaminated diet (100 or 500 µg CuCl_2_/g) while held for 12 days with free access to isosmotic seawater (25 ‰ salinity, 750 mOsm kg^-1^ H_2_O, 14 mmol L^-1^ Na^+^, 16 mmol L^-1^ Cl^-^). Data are the mean ± standard error (N= 10). *P≤ 0.05 compared to control crabs (0 µg CuCl_2_/g).

The gill Na^+^/K^+^-ATPase activity increased 1.3-fold in the crabs receiving the copper-contaminated diets (0.012 < P <0.023) compared to the control crabs (Figure 3). In contrast, gill carbonic anhydrase activity decreased by 19-35% in the copper-contaminated (0.004 < P < 0.037) compared to the control crabs (Figure 3).

**Fig. 3.**
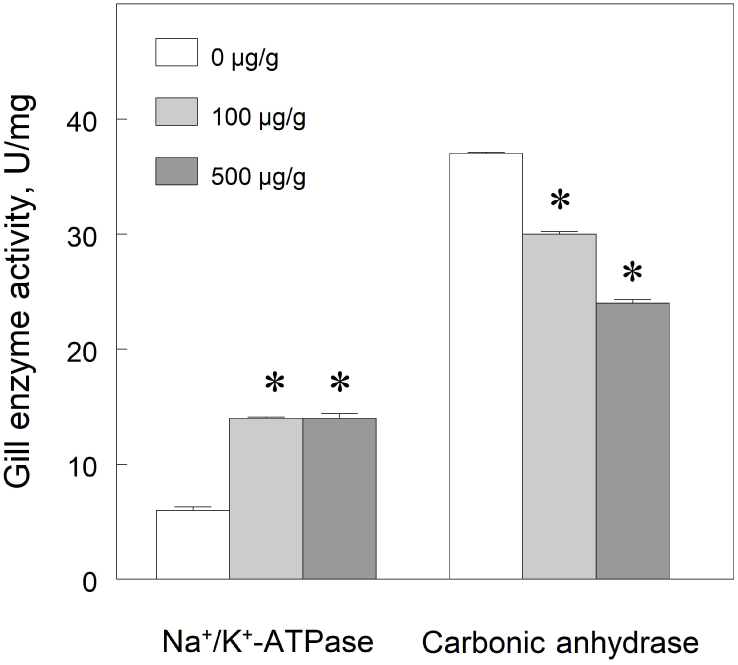
Na^+^/K^+^-ATPase activity and cytoplasmic carbonic anhydrase activity in homogenates of pooled anterior and posterior gills from *Minuca rapax* fed a copper-contaminated diet (100 or 500 µg CuCl_2_/g) for 12 days. Data are the mean ± standard error (N= 10). *P≤ 0.05 compared to control crabs (0 µg CuCl_2_/g).

The mechanisms of waterborne copper toxicity in crustaceans are starting to be better understood (Rainbow 2002, 2007; Capparelli et al., 2017). However, the uptake and toxicity of dietary-delivered metals have not been well investigated, particularly in semi-terrestrial fiddler crabs for which no data are available.

Our laboratory study of dietary copper delivery for 12 days in *Minuca rapax* showed that while elevated, copper concentrations of 100 and 500 µg/g were not lethal for the crabs, which did not avoid the contaminated diets. Measured copper bioaccumulation was highest in the hemolymph and hepatopancreas, and lowest in the gills in both contaminated and non-contaminated crabs. Interestingly, copper-contaminated crabs showed lower copper concentrations in the hemolymph and hepatopancreas than did control crabs, but higher gill concentrations.

Although delivered by a dietary rather than waterborne route, copper contamination did affect osmotic and ionic regulation in *M. rapax*. Hemolymph osmolality increased while sodium and chloride concentrations decreased. The increased activity of the gill Na^+^/K^+^-ATPase, the main ion-transporting enzyme, may have been induced by the reduced hemolymph Na^+^ and Cl^-^ concentrations, constituting a compensatory biochemical response. This same response pattern was seen in *M. rapax* subjected to waterborne copper (Capparelli *et al*. 2017). In contrast, gill carbonic anhydrase activity decreased, likely reflecting enzyme inhibition by copper accumulated in the gill tissue, as also seen during waterborne copper exposure (Capparelli *et al*. 2017). The effect on gill carbonic anhydrase suggests that acid-base balance and CO_2_ excretion also are affected by contamination via dietary copper. While waterborne copper delivered at just 50 µg/L disrupts many physiological and biochemical processes (Arnold 2005, Genz, J *et al*. 2011), *M. rapax* appears to possess effective mechanisms for dietary copper detoxification. The crabs are resistant to elevated copper concentrations, showing no mortality, suggesting that tissue specific copper concentrations did not reach lethal threshold levels consequent to dietary exposure at 500 µg Cu/g as used here. The reduced copper concentrations seen in the hemolymph and hepatopancreas of the copper-contaminated crabs suggest the activation of copper excretion mechanisms.

Considering the route of contamination and the mechanisms of dietary copper accumulation, higher concentrations and increases would be expected in the hepatopancreas and hemolymph rather than in the gills (Rainbow 2002, Ahearn *et al*. 2004). However, dietary copper can be absorbed and redistributed from the digestive system to the hemolymph and tissues where it accumulates as insoluble metal-rich granules derived from lysosomes as a result of metallothionein activity (Nassiri *et al*. 2000, Vogt and Quinitio 1994). The decreased copper concentrations seen in the hepatopancreas of *M. rapax* at 500 µg Cu/g and in the hemolymph at 100 and 500 µg Cu/g suggest copper excretion rather than its detoxification. Copper content increased in the gills of *M. rapax* only at 500 µg Cu/g, as also seen in fish (Shaw and Handy 2006, Handy 1996, Kamunde *et al*. 2001), possibly reflecting systemic copper redistribution via the hemolymph in these dietary contaminated crabs.

The shore crab *Carcinus maenas* exposed to sub-lethal or lethal waterborne copper (Rtal and Truchot 1996, Weeks *et al*. 1993) responds similarly to *M. rapax* while copper injected into the estuarine crab *Scylla serrata* is cleared from the hemolymph within 24 h. Bioaccumulation of lead in the blue crab *Callinectes danae* is greater in response to waterborne than dietary lead delivery (Bordón *et al*. 2018, 2019), likely owing to gill lead uptake as a consequence of osmoregulatory ion-uptake processes located in the gills.

The pattern of accumulation in dietary copper-contaminated *M. rapax* contrasts with that encountered in waterborne delivery studies (Capparelli *et al*. 2017) where the hemolymph and hepatopancreas showed higher titers compared to copper-free crabs. For semi-terrestrial crabs like *M. rapax* that inhabit burrows in muddy sand in estuarine mangrove environments, food source seems to be an important route of metal exposure. Nevertheless, copper-contaminated water flow over the gills and direct body contact may be as relevant as is exposure via particulate food sources.

With regard to effects on osmotic and ionic regulation, dietary copper increased hemolymph osmolality and decreased Na^+^ and Cl^-^ in *M. rapax* held at 25 ‰S, which may reflect toxic copper accumulation in the gills. *Minuca rapax* is an extremely euryhaline species (Thurman et al., 2017), is an excellent hyper/hypo-osmotic regulator (Capparelli *et al*. 2016), and is roughly isosmotic at 25 ‰S (780 mOsm kg^-1^ H_2_O). The increased osmolality seen in copper-contaminated crabs in an isosmotic salinity may reflect increased K^+^ and Ca^2+^ uptake and/or the presence of other osmolytes like NH_4_^+^ or free amino acids. Ammonia transport pathways are affected by copper contamination and elevated hemolymph ammonia correlates with increased copper in freshwater crayfish (Allinson *et al*. 2000). Copper exposure in *M. rapax* reduced hemolymph Na^+^ and Cl^-^ concentrations, suggesting impaired ion regulatory capability as seen in species contaminated by waterborne copper (Postel *et al*. 1998, Handy *et al*. 2003). Hemolymph Na^+^ and Cl^-^ concentrations are reduced in mercury-exposed crayfish (Wright and Welbourn 1993) and in copper- and cadmium-exposed amphipods, *Gammarus pulex* (Brooks and Mills 2003), owing to altered Na^+^/K^+^-ATPase activities.

The accumulation of copper in the gills suggests that while *M. rapax* can tolerate copper at concentrations up to 500 µg Cu/g, both acid-base equilibrium and osmoregulatory activities may be affected. Gill Na^+^/K^+^-ATPase activity increased in *M. rapax* fed a copper-contaminated diet in contrast to the decreased activity seen with waterborne contamination at concentrations above 100 µg Cu/L (Capparelli *et al*. 2017) and in response to contamination *in situ* (Capparelli *et al*. 2016). In *Carcinus maenas*, copper inhibits gill Na^+^/K^+^-ATPase activity leading to decreased sodium transport into the hemolymph (Handy *et al*. 2002). These effects derive from changes in enzyme conformation and/or in protein/lipid interactions (Henry *et al*. 2012). Just how dietary copper affects gill Na^+^/K^+^-ATPase activity is uncertain, although oxidative stress also increases concomitantly, and Na^+^/K^+^-ATPase activity is modulated by oxidative stress in fish (Hoyle 2007). However, few studies have examined ionic regulation in response to dietary copper contamination, an aspect that requires further study in fiddler and other semi-terrestrial crabs.

Gill carbonic anhydrase activity was inhibited in a concentration-dependent manner in *M. rapax* contaminated by dietary copper as seen in many crustaceans where copper is a potent inhibitor of carbonic anhydrase (Vitale *et al*. 1999). A similar disturbance of acid-base regulation by waterborne copper is apparent in *M. rapax* (Capparelli *et al*. 2017). Carbonic anhydrase activity in the gills of the estuarine crab *Neohelice granulata* is likewise inhibited (Bianchini *et al.*,2008), suggesting a role in copper-induced acid-base disturbance. Sub-lethal copper contamination can cause acid-base balance perturbation even at concentrations that fail to induce or result in only modest osmoregulatory disturbances (Wang *et al*. 1998). This inhibition suggests that acid-base regulation and CO_2_ excretion are affected by copper contamination via both dietary and waterborne exposure in *M. rapax.* Further studies on the effects of copper contamination on acid-base equilibrium and gas exchange in semi-terrestrial crabs are necessary, given that their frequent bimodal respiration in water and air must demand substantial physiological adjustments.

Most studies of copper toxicity have focused on aquatic organisms and gill contamination while investigations using dietary copper contamination center on digestive enzymes, fecundity and reproduction (De Schamphelaere and Janssen 2007, Kooijman 2000, Nogueira *et al*. 2004). However, our findings show that *M. rapax* accumulates dietary copper in the gills, and that transport and excretion mechanisms likely lead to decreased hemolymph and hepatopancreas copper titers. Further, dietary copper shows toxic sub-lethal effects, particularly on osmoregulatory processes such as lowered hemolymph Na^+^ and Cl^-^, and increased gill Na^+^/K^+^-ATPase and decreased carbonic anhydrase activities. Dietary copper does appear to act as an osmoregulatory stressor as seen with waterborne contamination, although a systematic analysis using dietary copper should be conducted. These findings are particularly relevant for semi-terrestrial crabs that spend long periods feeding on sediments, one of the main sources of metal contamination.

## Compliance with Ethical Standards

### Conflict of interest

The authors declare that they have no conflict of interest.

## Acknowledgments

This investigation was financed by the Fundação de Amparo à Pesquisa do Estado de São Paulo (FAPESP, 2011/22537-0 to JCM) from which MVC received doctoral scholarships (2011/08065-9 and 2013/10672-6). JCM received excellence in research scholarships from the Conselho Nacional de Desenvolvimento Científico e Tecnológico (CNPq 300662/2009-2, 303613/2017-3). MG is a Maytag Chair of Ichthyology. This study is part of a doctoral dissertation by MVC (Comparative Biology, FFCLRP/USP) and received support from the Coordenação de Aperfeiçoamento de Pessoal de Nível Superior (CAPES, 33002029031P8, finance code 001).

## Notes

### Competing Interest Statement

The authors have declared no competing interest.

